# XoxF and the Calvin-Benson Cycle Mediate Lanthanide-Dependent Growth on Methanol in *Bradyrhizobium* and *Sinorhizobium*

**DOI:** 10.1101/2025.06.30.662339

**Authors:** Charlotte R. Mineo, Jerry Jiang, N. Cecilia Martinez-Gomez

**Affiliations:** Department of Plant & Microbial Biology, University of California, Berkeley, Berkeley, California, USA

## Abstract

Nodule-forming bacteria play crucial roles in plant health and nutrition by providing fixed nitrogen to leguminous plants. Despite the importance of this relationship, how nodule-forming bacteria are affected by plant exudates and soil minerals is not fully characterized. Here, the effects of plant-derived methanol and lanthanide metals on the growth of nitrogen-fixing *Rhizobiales* are examined. Prior work has demonstrated that select *Bradyrhizobium* strains can assimilate methanol only in the presence of lanthanide metals; however, the pathway enabling assimilation remains unknown. In this study, we characterize *Bradyrhizobium diazoefficiens* USDA 110, *Bradyrhizobium sp*. USDA 3456, and *Sinorhizobium meliloti* 2011 to determine the pathways involved in methanol metabolism in previously characterized strains, other clades of *Bradyrhizobium*, and the more distantly related *Sinorhizobium*. Based on genomic analyses, we hypothesized that methanol assimilation in these organisms occurs via the lanthanide-dependent methanol dehydrogenase XoxF, followed by oxidation of formaldehyde via the glutathione-linked oxidation pathway, subsequent oxidation of formate via formate dehydrogenases, and finally assimilation of CO_2_ via the Calvin-Benson-Bassham (CBB) cycle. Transcriptomics revealed upregulation of the aforementioned pathways in *Bradyrhizobium sp.* USDA 3456 during growth on methanol. Enzymatic assays demonstrated increased activity of the glutathione-linked oxidation pathway and formate dehydrogenases in all strains during growth on methanol compared to succinate. ^13^C-labeling studies confirmed the presence of CBB intermediates and label incorporation during growth on methanol. Our findings provide multiple lines of evidence supporting the proposed XoxF-CBB pathway and, combined with genomic analyses, suggest that this metabolism is widespread among *Bradyrhizobium* and *Sinorhizobium* species.

**Importance:** Nitrogen-fixing soil bacteria such as *Bradyrhizobium* and *Sinorhizobium* promote plant growth while reducing dependence on artificial, energy-intensive fertilizers. Numerous studies have attempted to increase bacterial nitrogen fixation and colonization of plant tissues by identifying the micronutrients and plant exudates that promote successful symbiotic relationships. Among the compounds encountered by rhizobacteria, lanthanides have received little attention, despite reports that plant growth is affected by the presence of lanthanides. We characterized three agriculturally relevant *Bradyrhizobium* and *Sinorhizobium* strains, demonstrated that they gain the capacity to utilize methanol when lanthanides are present, and experimentally determined the pathway by which this metabolism occurs. This study provides a foundation for understanding the impacts of bacteria-mediated lanthanide metabolism on plant growth.

## Introduction

Nitrogen-fixing soil bacteria are crucial for global nitrogen cycles, high yield crop production, and reduced fertilizer needs (1). Microbes supply approximately 50% of the fixed nitrogen consumed by crops, with the rest coming from energy-intensive artificial fertilizers (2). Diverse bacteria have been shown to fix nitrogen both in free-living forms and in differentiated cells, which fix N_2_ within the low-oxygen environment present in specialized plant root organs called nodules (3). *Bradyrhizobium* and *Sinorhizobium* are two genera of nodule-forming bacteria that colonize a range of legume species in a strain-dependent manner, but are especially well-known for colonizing soybean and alfalfa, respectively (4).

*Bradyrhizobium*, *Sinorhizobium*, and other nodule-forming bacteria encounter diverse substrates as they migrate between the bulk soil and the carbon-rich rhizosphere, colonize root tissues, and fix N_2_ as bacteroids in root nodules (5). One such substrate is methanol, an abundant volatile carbon source in plant environments due to the activity of pectin methylesterases, which release methanol from both live plant tissues and plant detritus (6, 7). In above-ground plant tissues, methanol has been shown to drive colonization by allowing bacteria capable of using methanol as a sole source of carbon and energy, methylotrophs, to out-compete non-methylotrophic strains (8, 9). Similar mechanistic studies have not yet been conducted in soils or with N_2_-fixing bacteria, although it’s known that *Bradyrhizobium* assimilate methanol *in situ* in grassland soils and the rhizosphere (10).

Lanthanide (Ln) metals play a catalytic role in methanol metabolism in various plant-associated bacteria, including *Methylobacterium* and *Bradyrhizobium* (11–13). Lns are involved in methanol metabolism as cofactors in certain pyrroloquinoline quinone (PQQ) methanol dehydrogenases (PQQ-MDH) that oxidize methanol to formaldehyde. The Ln-dependent PQQ-MDH, XoxF, was first shown to be functional and require Lns in 2012 (11). To facilitate the binding of Lns, XoxF contains an additional Asp in the active site compared to the homologous, Ca-dependent PQQ-MDH, MxaF. This residue is essential for Ln-binding and catalysis and is used to predict Ln-dependency (14). Lns are introduced to soil through fertilizers (15–17) and can therefore influence the physiology of soil-dwelling microbes such as *Bradyrhizobium* and *Sinorhizobium*.

The activity of PQQ-MDHs, such as XoxF and MxaF, produces formaldehyde, an obligate, toxic intermediate that all methylotrophs must cope with during growth on methanol. Multiple pathways exist to oxidize formaldehyde and generate NAD(P)H, but the tetrahydromethanopterin (H_4_MPT) and glutathione (GSH)-linked oxidation pathways are most common (18). *Methylobacterium, Hyphomicrobium*, and many methanotrophs rely on the H_4_MPT pathway for formaldehyde oxidation (17, 18) while the GSH-linked pathway has been shown to play a supportive role in formaldehyde oxidation in *Methylobacterium aquaticum* 22A and *Rhodobacter sphaeroides* (19, 20), and is essential for growth on methanol in *Paracoccus denitrificans* (21). These pathways use different carbon carriers (H_4_MPT or glutathione), but similar reactions: both pathways immediately detoxify formaldehyde by covalently linking formaldehyde to their respective carbon-carrier, then oxidize the formyl group, and finally release the formyl-carbon from the carrier to recycle the carrier and generate formate. Formate can be further oxidized by formate dehydrogenases to generate CO_2_ and additional NAD(P)H (19). These pathways serve not only to detoxify formaldehyde and oxidize it for entry into assimilatory cycles, but also to generate NAD(P)H that fuels cellular processes and assimilatory cycles.

Carbon from methanol can be assimilated at several steps -formaldehyde, formate, or CO_2_-using diverse pathways, but the specific pathways employed by *Bradyrhizobium* and *Sinorhizobium* remain unknown. In Type I methylotrophy, Gamma and Betaproteobacteria use the ribulose monophosphate (RuMP) pathway to incorporate formaldehyde. Type II methylotrophs assimilate carbon at the level of formate. This group includes mostly Alphaproteobacteria, such as the model methylotroph *Methylobacterium extorquens* AM1. In *M. extorquens* AM1, approximately 50% of formate is oxidized to CO_2_ to generate NAD(P)H, the remaining 50% is incorporated into the serine cycle (20–22), and the ethylmalonyl-CoA (EMC) pathway is used to regenerate glyoxylate (23, 24).

As an alternative to the RuMP pathway and serine cycle, all carbon from methanol can be fully oxidized to CO_2_ and assimilated via the Calvin-Benson-Bassham (CBB) cycle. Operating the CBB cycle may appear inefficient due to the high energy requirements for CO_2_ fixation. However, use of the CBB cycle for growth on methanol has been validated in nitrogen-fixing Alphaproteobacteria such as *Beijerinckia mobilis* and *Xanthobacter flavus*, as well as the verrucomicrobial methanotroph *Methylacidiphilum fumariolicum* SolV, and the Gammaproteobacteria methanotroph *Methylococcus capsulatus* Bath (25–28). Keltjens et al. and Huang et al. have suggested that *Bradyrhizobium* and *Sinorhizobium* may be able to assimilate methanol using the CBB cycle, but this metabolism has yet to be experimentally validated (29, 30).

Despite their agricultural importance and ability to assimilate methanol *in situ* in grassland soils (10), very few studies have characterized methanol metabolism or Ln-dependency among the *Bradyrhizobium* and *Sinorhizobium.* In separate *in vitro* studies, *Bradyrhizobium diazoefficiens* USDA 110 and *Bradyrhizobium sp.* Ce-3 were shown to grow on methanol only when light Lns (lanthanum, cerium, praseodymium, or neodymium) were present, and growth was linked to the activity of XoxF (12, 13). To our knowledge, growth on methanol has not been reported among the *Sinorhizobium,* despite the *in vitro* characterization of a XoxF homolog from *Sinorhizobium meliloti* 5A14, which requires light Lns for activity (29). Notably, *Sinorhizobium meliloti* genes for methanol oxidation are located directly upstream of CBB cycle genes, such as RuBisCO, on the symbiotic megaplasmid pSymB (31). Growth on formate has been observed in *Sinorhizobium* and is known to be dependent on the presence of the CBB cycle and an NAD^+^-dependent molybdenum-containing formate dehydrogenase, Fds (32).

Based on bioinformatic analyses, we hypothesize that *Bradyrhizobium* and *Sinorhizobium* oxidize methanol using XoxF, detoxify formaldehyde via the GSH-linked oxidation pathway, produce CO_2_ using formate dehydrogenases (FDHs), and assimilate methanol-derived CO_2_ using the CBB cycle (Fig. 1) (hereafter referred to as the XoxF-CBB pathway). Transcriptomic, proteomic, metabolomic, and biochemical studies show that the XoxF-CBB pathway in *Bradyrhizobium sp.* USDA 3456 (*Bs.* 3456) is operational. Additionally, we use proteomics and biochemical assays to confirm that the XoxF-CBB pathway operates in another species, *Bradyrhizobium diazoefficiens* USDA 110 (*Bd.* 110), and the more distantly related *Sinorhizobium meliloti* 2011 (*Sm.* 2011). This is the first report of Ln-use and methanol assimilation by *Sinorhizobium*. Bioinformatic analyses suggest that methanol metabolism via the XoxF-CBB pathway is widespread among nitrogen-fixing *Rhizobiales*, including *Bradyrhizobium, Sinorhizobium,* and *Mesorhizobium*. This work reveals overlooked methanol metabolism pathways and new Ln-dependent species, allowing for a better understanding of the effects of methanol and Ln on nitrogen-fixing bacteria and agricultural settings.

**Fig 1.**
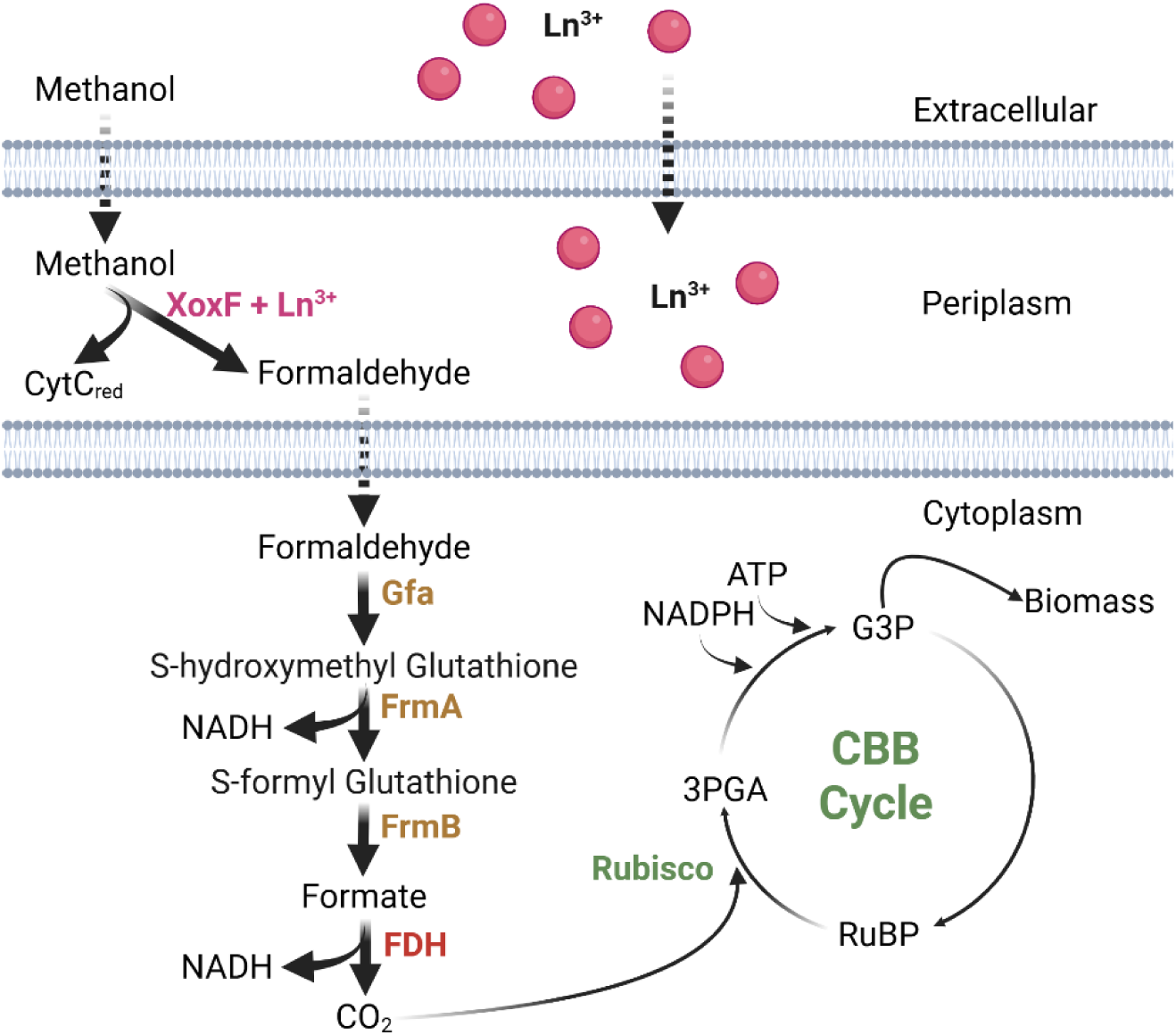
Schematic overview of the proposed XoxF-CBB Pathway for methanol metabolism in *Bradyrhizobium* and *Sinorhizobium*. La^3+^ is transported into the cell and complexes with apo-XoxF. XoxF (pink) oxidizes methanol to formaldehyde in the periplasm. Formaldehyde is oxidized to formate in the cytoplasm by the multi-step GSH-linked oxidation pathway (brown). Formate is oxidized to CO_2_ by an NAD^+^-dependent formate dehydrogenase (red). CO_2_ can be assimilated via the Calvin-Benson Cycle (green). For clarity, only the most relevant steps of the CBB Cycle are depicted. Dashed lines represent hypothesized steps based on similarity with other systems. Solid lines show experimentally supported steps from this study. XoxF - Lanthanide-dependent methanol dehydrogenase. Gfa - Glutathione-dependent formaldehyde activating enzyme. FrmA - glutathione-dependent formaldehyde dehydrogenase. FrmB - S- formyl glutathione dehydrogenase. FDH - NAD^+^-Dependent formate dehydrogenase. CBB Cycle - Calvin-Benson-Bassham Cycle. 3-PGA - 3-Phosphoglyceraldehyde. G3P - Glyceraldehyde 3 Phosphate. RuBP - Ribulose 1,5-bisPhosphate. Created in https://BioRender.com.

## Methods

### Media and Growth Conditions

All strains were grown at 30 °C with orbital shaking at 200 rpm in an Innova S44i incubator shaker (Eppendorf). Strains and plasmids used in this study are listed in Table 1. Pre-cultures of *Bs.* 3456, *Bd.* 110, *Sm.* 2011 and *Me* AM1 were grown in *Methylobacterium* PIPES (MP) defined minimal media (33) supplemented with either 1 g/L of gluconate, arabinose, and yeast extract or 15 mM succinate. When preparing growth curves or larger cultures for assays, pre-cultures were spun down at 2,000 x g for 10 minutes at 22 °C, washed in MP without carbon sources, and used to inoculate at a starting OD_600_ of 0.05-0.1. Growth was monitored at OD_600_ on either 50 mM methanol or 15 mM succinate with 10 μM LaCl_3_ when indicated. Experiments for Supplemental Figure 1 were conducted with the indicated Ln species in 650 μL cultures in transparent 48-well plates (Corning), incubated at 30 °C with orbital shaking at 548 rpm, with OD_600_ readings every 60 minutes using a Synergy HTX plate reader (Agilent). Otherwise, aerobic cultivation for growth curves occurred in sterile round-bottom 14 mL polypropylene tubes (Falcon). Experiments in varying oxygen conditions were performed in Balch-style tubes covered either with sterile tinfoil (unsealed) or plugged with blue stoppers (sealed condition). Growth was monitored by measuring OD_600_ using either the Ultrospec 10 (Amersham Biosciences) for polypropylene tubes or the Thermo Scientific GENESYS version 20 spectrophotometer for Balch-style tubes.

**Table 1.** Strains and plasmids used in this study.

| Strain or Plasmid | Description | Reference |
| --- | --- | --- |
| **Strain** |  |  |
| *Escherichia coli* 10β | Cloning – electrocompetent strain | Invitrogen |
| *Escherichia coli* S17 | Cloning – conjugation strain | New England BioLabs |
| *Methylobacterium extorquens* AM1 | Wild-type *Me.* AM1 | (58) |
| *Bradyrhizobium diazoefficiens* USDA 110 | Wild-type *Bd.* 110 | (59) |
| *Bradyrhizobium sp.* USDA 3456 | Wild-type *Bs.* 3456 | (60) |
| *Sinorhizobium meliloti* 2011 | Wild-type *Sm.* 2011 | (61) |
| *Methylobacterium extorquens* AM1 $\Delta mxaF$ | *Me.* AM1 $\Delta mxaF$ | (58) |
| *Bradyrhizobium diazoefficiens* USDA 110 $\Delta cbbS$ | *Bd.* 110 $\Delta cbbS$ | This Study |
| **Plasmids** |  |  |
| pCM433KanT | SacB allelic exchange backbone; Km<sup>R</sup> | (62) |
| pCRM031 | SacB allelic exchange donor *cbbS*; Km<sup>R</sup> | This Study |

### Biochemical Assays

50 mL cultures were grown in sterile 250 mL flasks with 10 μM LaCl_3_ and either 50 mM methanol or 15 mM succinate. All flasks were sealed with 45 mm gray bromobutyl rubber anaerobic flanges. Mid-log cultures (OD_600_ between 0.6 and 1) were immediately cooled on ice, centrifuged at 4 °C at 4,000 x g for 10 minutes, resuspended in 10 mL of 4 °C resuspension buffer (25 mM Tris, 10 mM NaCl, pH 8), centrifuged again at 4 °C at 4,000 x g for 10 minutes, and decanted. Cell pellets were stored at -80 °C for further analysis. Pellets were resuspended in 2 mL of resuspension buffer at 4 °C and lysed by a single pass at 25 kPSI through the MC-BA Cell Disruptor (Constant Systems). Lysed cells were centrifuged at 4 °C and 14,000 x g for 40 minutes in a Multifuge X Pro Series centrifuge (Thermo Scientific). All assays were performed at 30 °C in clear flat-bottom polystyrene 96-well plates (Greiner Bio-One). Absorbance, wavelengths indicated below, were monitored every 15 seconds for 5 minutes using a Synergy HTX plate reader (Agilent). A BCA kit (Pierce) was used to determine total protein content by measuring A_562_ in clarified lysates.

### MDH Activity Assay

MDH activity was assessed by monitoring the reduction of 2,6-dichlorophenolindophenol (DCPIP) according to the general procedures of Anthony and Zatman, but with modifications from Jahn et al. (34, 35). Briefly, 100 mM multi-component buffer at pH 9.0, and final concentrations of 1 mM phenazine ethosulfate (PES) and 100 μM DCPIP were used in the assay. Cell lysates were mixed at 4 °C with 185 mM ammonium chloride activator and then incubated for 1 minute at 30 °C before beginning the assay via the addition of assay mastermix containing either no substrate or 50 mM methanol. Absorbance was monitored at 600 nm and activity rates were calculated using an extinction coefficient of 19.7 mM^-1^ (35).

### GSH-Linked Formaldehyde Oxidation Assay

GSH-linked oxidation assays were performed according to the procedures of Yanpirat et al. (36) except that formaldehyde was prepared by heating and stirring a liquid solution of paraformaldehyde in the fume hood until the solution just began to boil, then immediately sealed. The production of NADH was monitored via absorbance at 340 nm using an extinction coefficient of 6.22 mM^-1^ to calculate activity (36).

### FDH Activity Assay

Protocols of Buttery and Chamberlain (37) were used to detect FDH activity by monitoring the reduction of *p*-iodonitrotetrazolium, except that no exogenous FDH was added; instead, cell lysates were used as the source of FDH. Cell lysates were incubated at 30 °C for 1 minute prior to the assay, and the reaction was initiated as previously described but using 1.67 mM formate. The reaction progress was monitored at 510 nm, and calculations of activity considered an extinction coefficient of 1.3 mM^-1^ for formazan (38).

### Transcriptomics in succinate with La and methanol with La

Cultures were grown as previously described, and samples were harvested when cultures reached mid-log (OD_600_ of 0.3 for methanol and 0.7 for succinate). Total RNA was extracted using the Qiagen RNeasy Kit as described by Okubo et al., except bead beating occurred for only 45 s and SUPERase•In was not needed (39). rRNA was removed via the Ribo-Zero RNA Plus rRNA Depletion Kit (Illumina), library prep, and sequencing were performed by SeqCoast Genomics, Portsmouth, NH. 12 million 2 x 150 bp paired-end reads were generated per sample. Paired-end libraries were uploaded to KBase (40), and reads were aligned to the *Bs.* 3456 genome (from MAGE accession 18212.1) with HISAT2 and assembled using StringTie and a PCA plot demonstrated separation between the samples grown on methanol and succinate (Fig. S1). DeSeq2 was used to analyze expression patterns. Genes were classified as differentially expressed using the stringent cut-off of Log_2_ Fold Change greater than 2 and -log_10_ q-value (p-value adjusted for the false discovery rate) greater than 7. Both conditions were analyzed in triplicate to generate Fig. 2.

**Fig 2.**
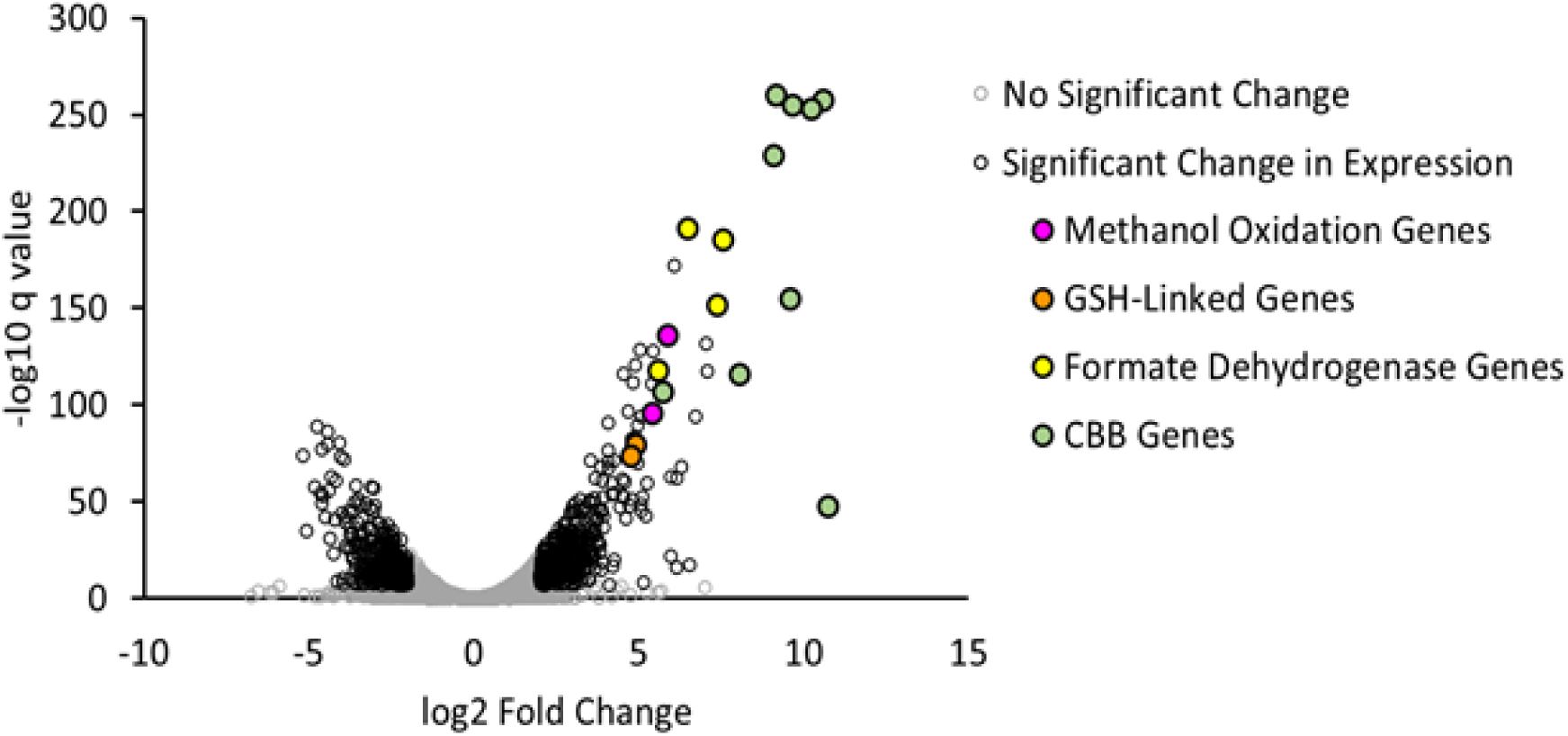
Volcano plot of differentially expressed genes during growth of *Bs.* 3456 on methanol and succinate. Cells were grown with either 15 mM succinate + La (left) or 50 mM methanol + La (right). Transcripts of interest for methanol metabolism are indicated by color-filled points. Genes were classified as differentially expressed using the stringent cut-off of Log_2_ Fold Change greater than 2 and -log_10_ q-value greater than 7. Using these parameters, 331 genes were downregulated, and 453 were upregulated out of a total of 10,757 unique genes detected via RNA-seq.

### SDS-PAGE and 1-D gel digestion proteomics

Clarified cell lysates were mixed with 2X Laemmli buffer + 5% beta-mercaptoethanol and boiled at 95 °C for 10 minutes. 20 μg of total protein was present in each sample. Samples were run on a 10-well, 4–20% gradient Mini-PROTEAN® TGX Stain-Free™ Gel at 150V with a PageRuler Prestained ladder (Thermo). The gel was stained using Coomassie Blue and imaged using the GelDoc Go, Software Version 3.0.0.07 ’Coomassie’ program (Bio-Rad). Single bands at the predicted molecular weight of RuBisCO (54 kDa) were excised, stored at -20 °C, and sent to the Vincent J Coates proteomics facility at UC Berkeley for 1-D reverse-phase LC-MS/MS.

### Plasmid and Strain Construction

A plasmid for the clean deletion of *cbbS* was constructed in the pCM433KanT backbone, which includes kanamycin-resistance genes for selection and *sacB* for counterselection with sucrose (20). Approximately 650 basepairs of genomic DNA directly upstream and downstream of *cbbS* (AAV28_09890) were amplified with overhangs (Table S1) from the *Bd.* 110 genome and inserted into the pCM433KanT plasmid using HiFi DNA Assembly Master Mix (NEB). The resulting plasmid was sequenced (Plasmidsarus) and electroporated into *Escherichia coli* S17. Biparental mating between *E. coli* S17 containing the deletion plasmid and wild-type *Bd.* 110 was used to introduce the plasmid. Mating took place for 48 hours on MP plates containing 1 g/L each of arabinose, gluconic acid, and yeast extract (MP-AGY plates). 5 μg/mL tetracycline and 100 μg/mL kanamycin were used for the first selection, followed by a transfer to 5 μg/mL tetracycline and 200 μg/mL kanamycin, both on MP-AGY plates. Kanamycin resistant single-crossovers were then transferred to MP-AGY plates supplemented with 10% sucrose, for selection, and 5 μg/mL tetracycline to inhibit the growth of contaminants, including *E. coli*. Resulting colonies were replica patched onto MP-AGY plates supplemented with either 10% sucrose or 200 μg/mL kanamycin. Those that grew on sucrose, but not kanamycin, were PCR screened for the loss of *cbbS* and sequenced (Plasmidsaurus) to confirm clean deletion of *cbbS*.

### Liquid Chromatography-Mass Spectrometry to detect 13-C labeled intermediates of the CBB cycle

Strains were pre-cultured in MP +10 μM LaCl_3_ with 50 mM unlabeled methanol as a sole carbon source. Mid-log pre-cultures (OD_600_ of 0.5) were centrifuged for 10 minutes at 2,000 x g, washed in MP without carbon sources, and transferred to 50 mL of MP media containing 50 mM of either 13-C labeled or unlabeled methanol (starting OD_600_ of 0.01). When preparing the cultures, each flask was sealed with gray bromobutyl flanges to promote assimilation of CO_2_ from methanol, instead of unlabeled CO_2_ from the atmosphere. These cultures were grown to mid-log (OD_600_ of 0.5) and harvested via vacuum filtration on 0.22 μm nylon filters (Micron Separators Inc). Filters were immediately flash-frozen in liquid nitrogen and were stored at -80°C until analysis.

A boiling water extraction was used according to the procedures of Cocuron and Alonso (41) except that tungsten beads were not needed to disrupt the tissues. Samples were flash frozen, lyophilized, and sent to the University of North Texas BioAnalytical Facility on dry ice. Samples were resuspended and analyzed via reverse-phase anion-exchange LC-MS/MS according to the procedures of Cocuron and Alonso 2014 (41). Analytes were detected via multiple reaction monitoring, and retention time and fragment masses were compared with external phosphoglyceraldehyde (PGA), fructose 6-phosphate (F6P), glucose 6-phosphate (G6P), ribulose 5-phosphate (R5P), ribulose 1,5-bisphosphate (RuBP), and sedoheptulose 7-phosphate (S7P) standards. The total peak area for each mass isotopomer was calculated and used for comparisons between samples.

### Structural predictions and determination of homology between putative XoxFs

The amino acid sequence of XoxF1 from *Me. AM1* (META_1740) was queried against the genomes of *Me.* AM1 (chromosome META1.1), *Bd*. 110 (chromosome CP011360.1), *Bs.* 3456 (WGS VIDU01.1), and *Sm.* 2011 (plasmid NC_020560.1, plasmid NC_020527.1, chromosome NC_020528.1) using the BLAST functionality in MAGE, MAgnifying Genomes (42). 21 hits were returned from among the four genomes, and all but one were annotated as PQQ dehydrogenases. A manual search of amino acid sequences revealed the presence/absence of an additional aspartate, which was shown to be essential for Ln-binding (14). Sequences from the BLAST search were aligned using Clustal W, a phylogenetic tree was constructed using IQTree, and the tree was visualized using iTOL (43–45). All sequences were fed to AlphaFold3 (46) using default parameters, and the overall fold without ligands was assessed. AlphaFold3 models of key PQQ-DH homologs were aligned with the crystal structure of the XoxF1 holoenzyme from *Me.* AM1 (PDB ID: 6OC6) using ChimeraX (47).

### Synteny visualization using CLINKER

Organisms were selected based on literature reports of relevant 1-carbon metabolism and to provide a diverse array of *Rhizobiales*. Sequences were manually identified based on proximity to XoxF homologs and downloaded from the IMG database and aligned via CLINKER (online) with default settings (48, 49).

### Predicting the distribution of the XoxF-CBB pathway among *Rhizobiales* genomes

Select *Rhizobiales* genomes were accessed and analyzed using the IMG database (49). Genomes were searched by BRENDA enzyme commission (EC) numbers to avoid missing homologs due to varied nomenclatures. Methanol oxidation genes were Ca-dependent MDH (MxaF, 1.1.2.7) and Ln-dependent MDH (XoxF, 1.1.2.10). GSH-linked oxidation genes were S-(hydroxymethyl) glutathione synthase (Gfa, 4.4.1.22), S-(hydroxymethyl) glutathione dehydrogenase (FrmA, 1.1.1.284), S-formylglutathione hydrolase (FrmB, 3.1.2.12). Select genes of the H_4_MPT pathway were formaldehyde activating enzyme (Fae, 4.2.1.147) and methenyltetrahydromethanopterin cyclohydrolase (Mch, <u>3.5.4.27</u>). Genes of the H_4_F pathways were formyltetrahydrofolate deformylase (PurU, 3.5.1.10), and formate-tetrahydrofolate ligase (FtfL, 6.3.4.3). Formate dehydrogenase was (FDH, 1.17.1.9). 3-hexulose-6-phosphate synthase (Hps, <u>4.1.2.43</u>) and 3-phospho-6-hexuloisomerase (Phi, <u>5.3.1.27</u>) were used as markers for the RuMP cycle. The Glyoxylate Shunt was identified by isocitrate lyase (Icl, 4.1.3.1) and malate synthase (<u>2.3.3.9</u>). The CBB was searched using RuBisCO (CbbL, 4.1.1.39) and phosphoribulokinase (Prk, 2.7.1.19). The serine cycle was predicted from glycine hydroxymethyltransferase (GlyA, 2.1.2.1), malate thiokinase (Mtk, 6.2.1.9), and malate-CoA lyase (Mcl, 4.1.3.24). The EMC pathway was indicated by crotonyl-CoA carboxylase/reductase (CcR, 1.3.1.85) and (2S)-methylsuccinyl-CoA dehydrogenase (<u>1.3.8.12</u>). The phylogenetic tree was constructed from RpoB amino acid sequences extracted from the IMG database and analyzed using ClustalW, IQTree, and visualized using iTOL. The tree was manually annotated to reflect gene presence/absence according to the above analysis.

## Results

### Genomic data suggests the CBB cycle is employed for methanol assimilation

Because multiple methanol assimilation modules have been characterized in the literature, we initially explored genomic data to identify potential pathways employed by strains *Bs.* 3456, *Bd.* 110, and *Sm.* 2011. All three strains lack key genes for both the serine cycle (malate thiokinase, malyl-CoA ligase) and RuMP pathway (3-hexulose-6-phosphate synthase, 3-phospho-6-hexuloisomerase) (Table S2), and the lack of these genes indicates that the serine and RuMP cycles cannot function in methanol assimilation in *Bs.* 3456, *Bd*. 110, and *Sm*. 2011. Other serine cycle and RuMP pathway genes that are present in these strains likely function in overlapping metabolic pathways, such as the citric acid cycle or the pentose phosphate pathway. However, all three strains possess genes predicted to encode the Ln-dependent dehydrogenase XoxF, the glutathione-linked oxidation pathway, various FDHs, and a complete CBB cycle (Table S3). This suite of genes led us to hypothesize that strains *Bs.* 3456, *Bd.* 110, and *Sm.* 2011 can oxidize methanol to CO_2_ and assimilate this carbon using the CBB cycle (Fig. 1).

The presence of *xoxF* homologs led us to search for appropriate accessory genes for Ln uptake in the organisms of interest. Homologs for genes of the *lut* (lanthanide utilization and transport) cluster from *Me.* AM1, including *lutA*, *B*, *E*, *F*, and *G*, encoding an ABC transport system and periplasmic proteins, can be found in all strains with 40-60% identity, according to the BLAST function in MAGE. Homologs of *lutH*, the TonB-dependent outer membrane transporter, from *Bd*. 110 and *Bs.* 3456 exhibit lower similarity to the sequence from *Me.* AM1 than the other *lut* genes, and no obvious homolog can be found in *Sm.* 2011 using BLAST (Table S3). While it’s possible that a *lut* cluster operates in these organisms, it is unlikely that they utilize the small molecule Ln-chelator methylolanthanin for Ln solubilization, as none of the strains of interest encode clear homologs of *mllA-E* according to BLAST analysis in the MAGE database (50).

### Transcriptomics reveal upregulation of the proposed methanol assimilation genes

To identify genes likely to be involved in methanol metabolism, RNA-seq was performed on *Bs.* 3456 grown either on 15 mM succinate or 50 mM methanol as sole carbon sources, supplemented with 10 μM La in both conditions. When stringent cut-offs of log_2_ fold change greater than 2 and -log_10_ q-value greater than 7 were applied, 453 genes were identified as upregulated in the methanol condition compared to the succinate condition. Several genes necessary for methanol metabolism were among the most highly upregulated including: *xoxF* (VIDU01_860227), putative *xoxG* (VIDU01_860228), *gfa* (VIDU01_860231), *frmA* (VIDU01_860229), genes for NAD-dependent formate dehydrogenase subunits alpha, beta, delta, and gamma (VIDU01_10810, 10811, 10808, and 10812, respectively), *cbbL* and *cbbS* (VIDU01_10123 and 10124), and other genes of the CBB cycle (Fig. 2). Genes potentially involved in the serine/EMC cycle or the RuMP pathway are either downregulated or do not change in expression in *Bs.* 3456 during growth on methanol vs succinate (Log_2_ Fold Change of 1.5 or less). This includes serine cycle and EMC pathway genes such as *glyA* (VIDU01_310356), *ppc* (VIDU01_10600), *mdh* (VIDU01_270073), *ccr* (VIDU01_510004), as well as *icl* (VIDU01_320092) and *ms* (VIDU01_640301) from the glyoxylate shunt.

### Identification of XoxFs driving light Ln-dependent methanol oxidation

The ability to assimilate methanol was expected for *Bd.* 110 due to prior reports in the literature (13). Here, we show that *Bd.* 110, *Bs.* 3456 and *Sm.* 2011 grow on methanol only when light Lns are present (Fig. 3A, Fig. S2). Higher concentrations of Lns support faster growth rates, with a maximum growth rate occurring at 1-10 μM La. (Fig. S2). In addition to the presence of Lns, *Bd.* 110 required sealed Balch-style tubes for growth on methanol (Fig. S3).

**Fig 3.**
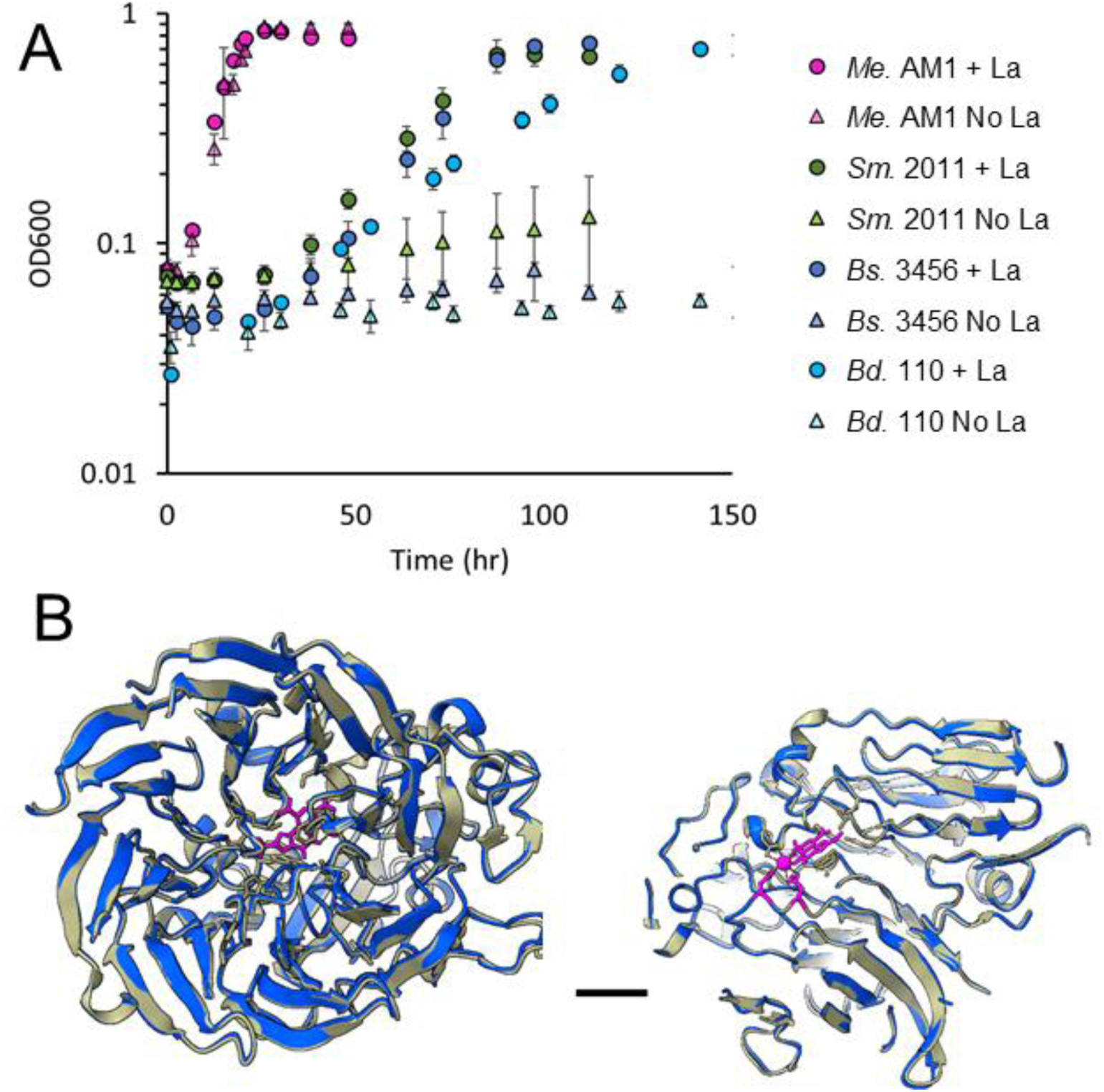
Phenotypic and structural evidence for XoxF as a methanol dehydrogenase involved in Ln-dependent growth on methanol. **(A)** Growth of wild-type *Me.* AM1 (pink), *Sm.* 2011 (green), *Bs.* 3456 (dark blue), and *Bd.* 110 (light blue) on 50 mM methanol with (circles) and without (triangles) 10 μM LaCl_3_. No significant differences in growth rate were found for *Me.* AM1 ± La. p≤0.001 ± La according to a 2-tailed t-test. Differences were significant ± La for strains *Bs.* 3456, *Bd.* 110 and *Sm.* 2011. **(B)** Comparison of overall structure and Ln-binding residues between the *Bs.* 3456 XoxF (VIDU01_860227) modeled with >90% confidence in Alpha Fold3 (blue) and the crystal structure of XoxF1 from *Me.* AM1 (tan) with PQQ and Ce present (PDB 6OC6). PQQ, Ce, and the catalytic and Ln-binding Asps from both models are highlighted in pink. Scale bar = 10Å.

Potential XoxF homologs from *Bs.* 3456, *Bd.* 110, and *Sm.* 2011 were compared to the crystal structure of XoxF1 from *Me.* AM1 (PDB structure 6OC6). *Me.* AM1 encodes 3 different Ln-dependent dehydrogenases, but XoxF1 is the principal PQQ-MDH for methanol oxidation (14). Similarly, *Bd.* 110, *Bs.* 3456, and *Sm.* 2011 each encode multiple predicted Ln-dependent PQQ-dehydrogenases (Fig. S4). We identified 1 homolog from each organism that has the highest sequence similarity to XoxF1 from *Me* AM1. Notably for *Bs.* 3456, the closest XoxF homolog (VIDU01_860227) is also the most highly upregulated PQQ dehydrogenase during growth with methanol and La compared to succinate and La (Fig. 2). All XoxF homologs were 601-602 amino acids in length and shared approximately 75% identity with XoxF1 from *Me*. AM1. When modeled in AlphaFold 3, all structures were predicted with >90% confidence except for an approximately 25 amino acid sequence at the N-terminus that is predicted to be a signaling peptide that promotes trafficking to the periplasmic space, according to InterPro (51). The presence and orientation of residues in the active site is particularly important for substrate specificity and for the ability to bind Ln^3+^. A model of VIDU01_860227 is shown compared to XoxF1 from *Me.* AM1, and a magnified view highlights the conserved orientation of the predicted catalytic aspartate and the Ln-binding aspartate (pink) (Fig. 3B). The putative XoxFs from *Bd.* 110 and *Sm.* 2011 adopt a similar overall fold and active site conformation.

### GSH-linked genes are adjacent to *xoxF*, and the GSH-linked pathway for formaldehyde oxidation is active during growth on methanol

Based on genomic evidence from all strains and transcriptomic evidence from *Bs.* 3456, we predicted that the GSH-linked oxidation pathway was used to oxidize formaldehyde to formate during growth on methanol (Fig. S2 and Fig. 2). In addition, synteny analysis shows components of the GSH-linked oxidation pathway (*gfa*, *frmA*, *frmB*) directly adjacent to XoxF homologs in *Sm*. 2011, *Bd.* 110, and *Bs.* 3456 (Fig. 4A). Similar organization is observed in select *Paracoccus denitrificans* and *Rhodobacter sphaeroides,* and *Sinorhizobium medicae* strains. In organisms that do not use the CBB, like *Me.* AM1 and *Xanthobacter autotrophicus*, *mtdA* and *fch* are involved in formaldehyde metabolism and are adjacent to *xoxF* homologs in their respective organisms. This conserved organization suggests an evolutionary pressure to co-regulate these genes across divergent organisms and pathways.

**Fig 4.**
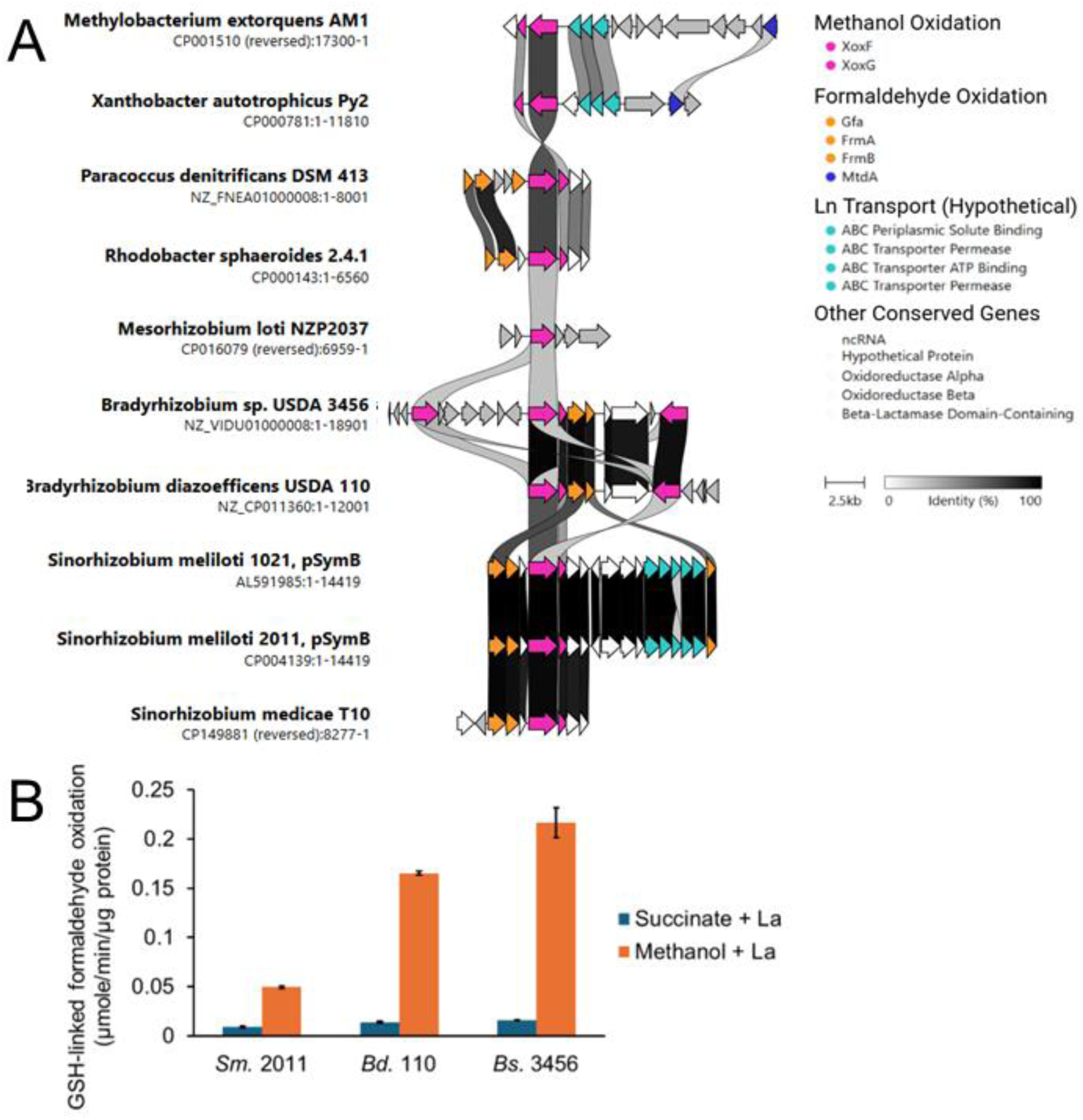
GSH-linked pathway implicated in methanol metabolism. **(A)** Synteny analysis of the region surrounding XoxF homologs in select Alphaproteobacteria. Arrows represent genes. Colored arrows are homologs that have greater than 30% sequence similarity with other selected strains. Gray arrows have less than 30% similarity to other genes in this dataset or have no homologs. White arrows are homologs with no known role in methanol metabolism. Homologous genes are connected by ribbons and darker ribbon color indicates higher similarity. **(B)** NADH-linked activity assay for glutathione-dependent formaldehyde dehydrogenase activity. All strains exhibit a statistically significant (p ≤0.001) difference in activity between succinate and methanol grown cultures according to a 2-tailed student’s t-test.

We used the GSH-linked oxidation assay, which monitors NADH produced by the GSH-dependent formaldehyde dehydrogenase, FrmA, to demonstrate that the GSH-dependent formaldehyde oxidation pathway is functional during growth on methanol. Lysates from *Sm.* 2011, *Bs.* 3456, and *Bd.* 110 cells grown on methanol have 5-, 11-, and 13-times greater GSH-dependent formaldehyde oxidation activity, respectively, compared to lysates from cells grown on succinate (Fig. 4B). Notably, activity from *Sm.* 2011 extracts grown on methanol is only 25% of the activity observed with *Bs.* 3456 extracts grown on methanol.

### NAD^+^-dependent FDH activity is elevated during growth on methanol

FDH activity is needed to generate NADH and produce CO_2_ for carbon assimilation via the CBB cycle. To determine whether FDHs were more active in cells grown on methanol versus succinate, we employed a formate dehydrogenase activity assay to quantify the ability of cell lysates to convert formate to CO_2_. *Me.* AM1 encodes four FDHs with redundant activity (19) and one or more homologs of each FDH type is found in the *Bd.* 110 and *Bs.* 3456 genomes (Table S3). In contrast, a single NAD^+^-dependent homolog of Fdh2A was identified in *Sm.* 2011 (Table S3). As expected, for all strains, there was greater FDH activity in lysates from cells grown on methanol than those grown on succinate (Fig. 5). Activity in methanol-grown cultures, compared to succinate-grown cultures, was 8-fold greater for *Bd.* 110 and 16-fold greater for *Bs.* 3456. FDH activity observed from *Sm.* 2011 lysates grown on methanol was less than 10% of that observed for *Bs.* 3456, but activity from *Sm.* 2011 cultures was 2-fold higher in methanol-grown cultures than succinate-grown cultures.

**Fig 5.**
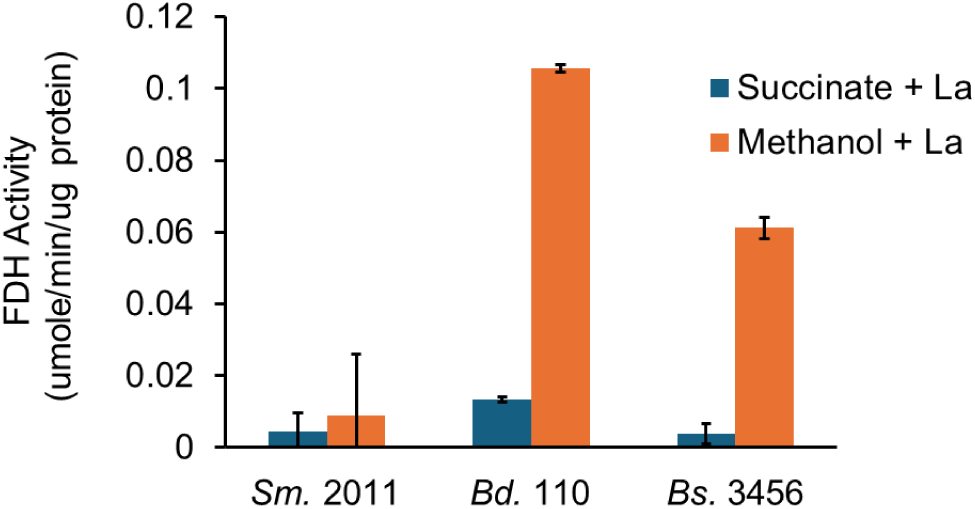
NAD^+^-dependent formate dehydrogenase activity assay. All strains exhibit a statistically significant (p ≤0.005) difference in activity between succinate and methanol grown cultures according to a 2-tailed student’s t-test.

### The CBB cycle is used for assimilation during growth on methanol

Formate oxidation by FDHs generates CO_2_, which we hypothesized is assimilated via the CBB cycle. Our transcriptomics from *Bs.* 3456 indicate that the CBB cycle is upregulated during growth on methanol compared to succinate (Fig. 2). To validate a functional CBB cycle, the presence of the large subunit of RuBisCO (CbbL) was confirmed in all our organisms of interest during growth on methanol via mass spectrometry by analyzing the 55-70 kDa region from the SDS-PAGE gel for *Bd.* 110, and *Bs.* 3456 samples grown on methanol and on the 40-55 kDa region for *Sm*. 2011 (Fig. 6). A Type IC RuBisCO was detected in all of these samples with 28-49% coverage of the CbbL sequence. Unlike the *Bradyrhizobium*, *Sm.* 2011 encodes both a Type IC RuBisCO (SM2011_b20198) and a Type II RuBisCO (SM2011_b20393), but only the Type IC was detected in this data set.

**Fig 6.**
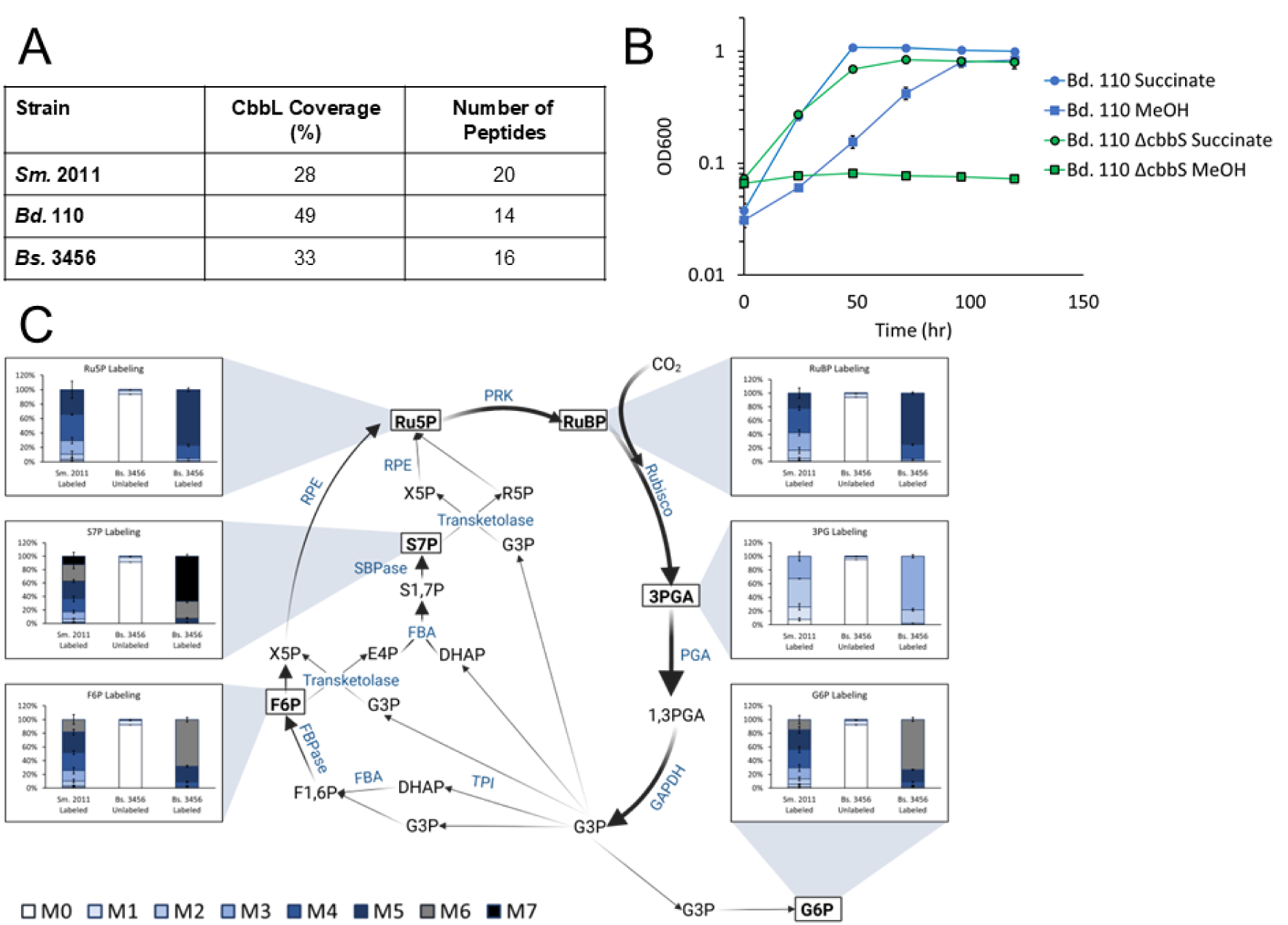
Involvement of the CBB cycle in methanol metabolism**. (A)** Coverage and number of peptides from Rubisco detected in methanol-grown lysates of *Bd.* 110, *Bs.* 3456, and *Sm.* 2011 via gel digestion LC-MS. **(B)** Growth of wild type (blue) and Δ*cbbS Bs.* 110 (green) on 15 mM succinate + 10 μM La (circles) and 50 mM methanol + 10 μM La (squares) in sealed Balch-style tubes. **(C)** The percentage of each mass isotopomer for 6 different analytes is depicted for labeled *Sm.* 2011, unlabeled *Bs.* 3456, and labeled *Bs.* 3456 with standard deviations. The isotopomer color-coding key at the bottom holds true for all analytes. Metabolites are spatially organized according to their role in the canonical CBB cycle and relative isotopomer abundance is shown near each measured metabolite. Data is the result of 3 biological replicates analyzed via reverse-phase anion-exchange LC-MS utilizing multiple reaction monitoring. Metabolites are given in black, and enzymes are shown in blue. Metabolites with a box around them have previously been detected via this method, and were analyzed in this experiment. Arrow thickness represents predicted approximate flux through each step of the pathway according to standard models of the CBB.

The essentiality of RuBisCO, and by extension the CBB cycle, for growth of *Bd.* 110 on methanol was also confirmed genetically by constructing a clean deletion of *cbbS* in *Bd.* 110. *Bd.* 110 Δ*cbbS* was unable to grow on methanol across three biological replicates (Fig. 6B). *Bd.* 110 Δ*cbbS* was still able to grow on succinate at levels similar to wildtype, indicating that lack of growth on methanol is due to the assimilatory role of RuBisCO, and not due to non-specific deficits of this strain. Due to challenges with molecular cloning in strains *Bs.* 3456 and *Sm.* 2011, deletions of *cbbS* could not be generated and alternative approaches were employed to provide evidence for operation of the CBB cycle in these strains.

*Bs.* 3456 and *Sm.* 2011 were fed 13C-methanol and incorporation of labeled carbon into CBB cycle intermediates was used for confirmation of CBB cycle activity. All flasks were sealed during growth on ^13^C methanol to prevent the escape of ^13^CO_2_, and cultures of *Bs.* 3456 and *Sm.* 2011 fed with ^13^C methanol underwent multiple doublings, 2.6 for *Sm.* 2011 and 6.2 for *Bs.* 3456, leading to a high degree of ^13^C labeling in the analytes of interest. All CBB cycle metabolites of interest (R5P, RuBP, 3PGA, G6P, F6P and S7P) were successfully detected and labeled in cultures of *Bs.* 3456 and *Sm.* 2011 fed ^13^C methanol (Fig 6A). For *Bs.* 3456 and *Sm.* 2011 samples, less than 2% of the M0 mass isotopomer remained for all analytes indicating a high degree of label incorporation. For *Bs.* 3456, the heaviest possible isotopomer was the most prevalent for each analyte whereas the second heaviest was most prevalent for *Sm.* 2011, consistent with fewer doublings from *Sm.* 2011. Unlabeled controls from *Bs.* 3456 cultures fed ^12^C methanol exhibited labeling patterns consistent with the isotopic distribution of unenriched carbon and with the observed isotopomer distributions of the unlabeled external standards.

### Genomic analyses suggest that the capacity for XoxF-CBB metabolism is prevalent among *Bradyrhizobium, Mesorhizobium,* and *Sinorhizobium*

To address whether the XoxF-CBB pathway is widespread within Alphaproteobacteria, diverse members of the *Rhizobiales* clade were queried for the presence of methanol assimilation genes using EC number searches in the IMG database. As a control, genomes of *Methylobacterium* strains were analyzed using the same methodology and the four *Methylobacterium* strains tested encode genes of the H_4_MPT pathway, serine cycle and EMC pathway. All tested members of the *Mesorhizobium*, *Sinorhizobium*, and *Bradyrhizobium* clades lack key genes for the serine/EMC pathway. None of the selected strains encode a homolog of hexulose-6 phosphate synthase, and only *Mesorhizobium sp.* NZP2037 encodes a copy of 6-phospho 3-hexuloisomerase for operation of the RuMP pathway (data not shown). However, many of these strains are predicted to encode genes for XoxF, the GSH-linked pathway, FDHs, and the CBB cycle (Fig. 7).

**Fig 7.**
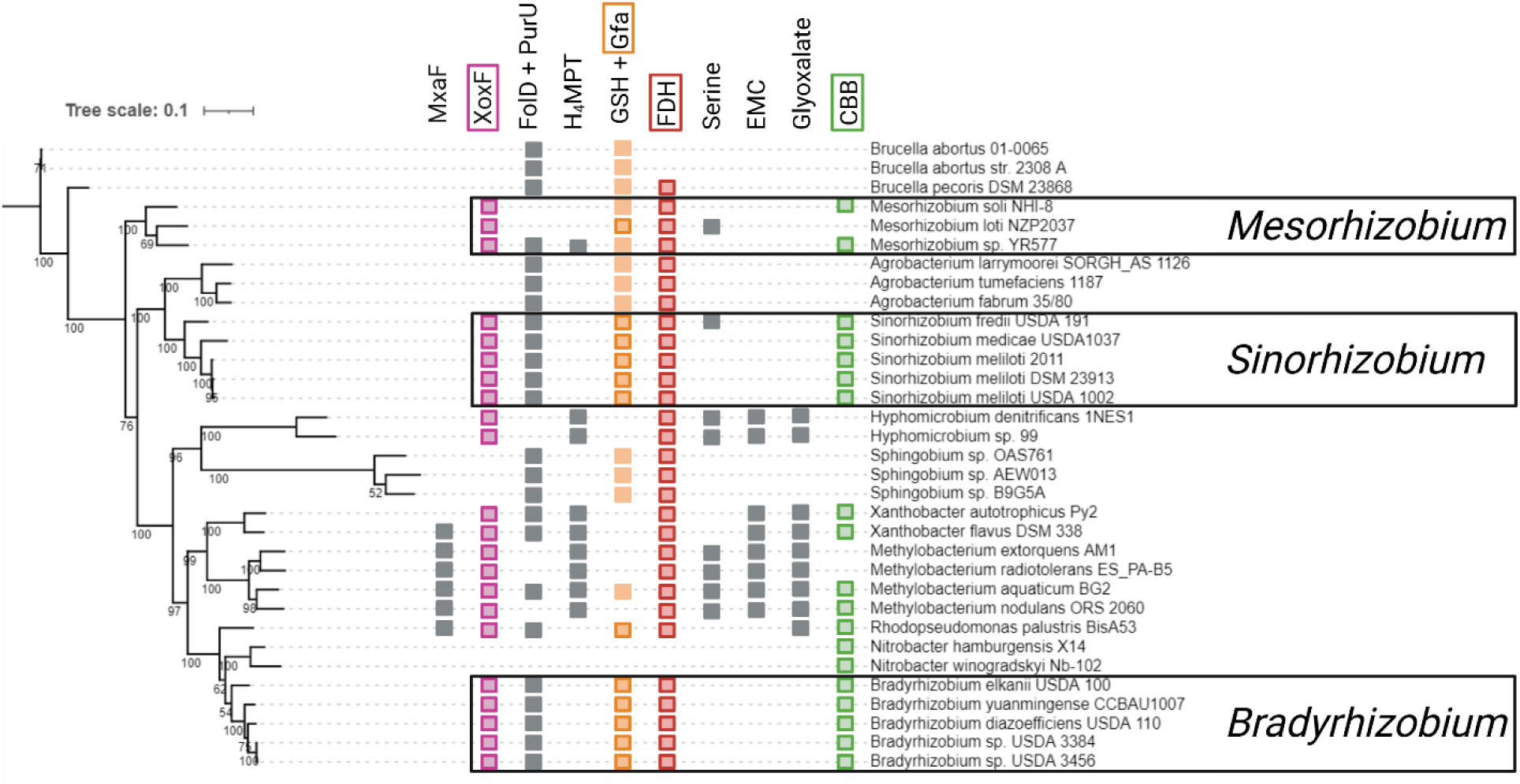
Genomic analyses suggest that the XoxF-CBB pathway for methanol metabolism is widespread among the *Rhizobiales*. Representative *Rhizobiales* were randomly selected and the phylogenetic tree was constructed using amino acid sequences of RpoB with bootstrap values given at each node. The tree was manually annotated to depict the presence/absence of methanol-utilization genes according to searches by EC number in the IMG database. Squares represent the presence of select genes and pathways; no square indicates that no homolog could be found. Steps of the XoxF-CBB pathway are indicated using colored squares and gray squares are used for other potential methanol assimilation pathways. Created in https://BioRender.com.

## Discussion

Lns are being introduced, both intentionally and inadvertently, at variable concentrations into agricultural soils via commercial fertilizers, making it important to understand how these metals might influence the growth of microbes and, consequently, the productivity of crops (52). The role of Lns in methanol metabolism of the model methylotroph *M. extorquens* has been characterized *in planta* and extensively characterized *in vitro* (14, 50, 53, 54). Lns have been shown to enable growth on methanol in *Bradyrhizobium*, but the pathway enabling growth on methanol and the extent of this metabolism among nitrogen-fixing *Rhizobiales* has yet to be determined (12, 13) despite their essential roles as rhizosphere plant-symbionts. Here, transcriptomic, biochemical, metabolomic, and bioinformatic evidence demonstrate that the characterized *Bradyrhizobium* and *Sinorhizobium* strains oxidize methanol using the Ln-dependent PQQ-MDH XoxF, oxidize formaldehyde using the GSH-linked oxidation pathway, oxidize formate via FDHs, and assimilate carbon as CO_2_ using the CBB cycle.

Evidence for the complete XoxF-CBB pathway is provided by transcriptomics of *Bs.* 3456 in which all of the modules involved in methanol assimilation were upregulated during growth on methanol compared to succinate (Fig. 2). Because La was present in both conditions, there was no upregulation of the predicted Ln transport and uptake cluster. Similarly, the lack of upregulation of the PQQ biosynthesis genes, suggests that PQQ is synthesized regardless of carbon source, or that the presence of Lns leads to the expression of PQQ biosynthesis genes in both conditions. XoxF1 (VIDU01_860227) is predicted to be Ln-dependent, has 76% similarity to XoxF1 from *Me.* AM1, and is the most highly upregulated methanol dehydrogenase, suggesting either a key role in methanol metabolism or regulation by substrate availability.

Consistent with the upregulation of a *xoxF*1 homolog, growth on methanol is completely dependent on the presence of Ln-metals for wild type *Bs.* 3456 (Fig. 3A). Additionally, growth rate is ‘tunable’ and increases as the amount of La in the media increases up to a maximum at 1-10 μM (Fig. S2). Seven other PQQ dehydrogenases are encoded by *Bs.* 3456 with lower similarity to XoxF1 from *Me.* AM1, but their functions are currently unknown (Fig. S4). Assays for FrmA and FDH activity, together with ^13^C labeling, confirm that the GSH-linked oxidation pathway, FDHs, and CBB cycle are functional during growth on methanol, consistent with the transcriptomic data. Detection of labeled intermediates from cultures supplied with ^13^C methanol further indicate that *Bs.* 3456, and *Sm.* 2011, are capable of assimilating methanol-derived ^13^CO_2_, and are not reliant on atmospheric ^12^CO_2_ (Fig. 6).

In our hands, *Bd.* 110 was unable to grow on methanol and La unless cultured in sealed Balch-style tubes (Fig. S3). Our initial attempts to grow *Bd.* 110 on methanol and La were unsuccessful and we hypothesized that *Bd.* 110 could be sensitive to oxygen due to the poor selectivity of RuBisCO for CO_2_ over O_2_. We sealed Balch-style tubes, such that the microbes have a finite amount of oxygen available, and compared to unsealed tubes where diffusion maintains atmospheric oxygen levels. Oxygen cannot be fully removed, due to the requirement for a strong electron acceptor for methanol oxidation in most organisms. *Bs.* 3456 exhibited a shorter lag in sealed tubes compared to foil-covered tubes, but *Bd.* 110 was only able to grow in the sealed tubes (Fig S2). *Me.* AM1, which does not use the CBB cycle, exhibits a slightly reduced growth rate in sealed tubes compared to foil-covered tubes. While other O_2_ or CO_2_-sensitive enzymes besides RuBisCO could be responsible for these results, the striking requirement for sealed conditions for the growth of *Bd.* 110 on methanol indicates that screens for Ln-dependent methylotrophs need to incorporate a low-oxygen condition, and this may be especially important for organisms predicted to use the CBB cycle.

RuBisCO has been shown to be important in multiple plant-*Bradyrhizobium* interactions, though the exact mechanisms for this effect remain unclear. A photosynthetic strain, *Bradyrhizobium sp.* ORS278, in symbiosis with *Aeschynomene indica,* exhibits a defect in symbiotic nitrogen fixation, but no defect in free-living nitrogen fixation when *cbbL* was disrupted (55). Notably for our study, when the *cbbL* sequence in *Bradyrhizobium diazoefficiens* USDA 110 (*Bd.* 110 in our shorthand) is disrupted, the mutants are less able to colonize roots and occupy nodules of soybean, suggesting that RuBisCO is involved in establishing plant-microbe symbioses (56). The role of RuBisCO in methanol metabolism for these *Bradyrhizobium* strains may contribute to these phenotypes, but further work is needed to confirm this hypothesis.

To our knowledge, the Ln-dependent growth on methanol observed here (Fig. 3) is the first report of Ln-dependent metabolism among the *Sinorhizobium*. Growth on methanol requires nanomolar to micromolar concentrations of La, and only light lanthanides facilitate the growth of *Sm.* 2011 on methanol (Fig. S2). In *Sm.* 2011, genes for XoxF, the GSH-linked pathway, Ln uptake, the CBB cycle and PQQ biosynthesis are found adjacent to one another in a 28.5 kb region on the symbiotic megaplasmid pSymB. Proteomic analysis of gel-digested bands indicated that *Sm.* 2011 expressed the Type Ic RuBisCO found in this cluster, instead of the Type II RuBisCO that is encoded further downstream on pSymB. Having nearly the complete suite of genes implicated in XoxF-CBB metabolism found on a megaplasmid increases the likelihood of horizontal gene transfer for methanol-related genes among the *Sinorhizobium* (57).

Like the *Bradyrhizobium*, *Sm.* 2011 cultures grown on methanol exhibit greater GSH-linked oxidation activity than those grown on succinate. However, *Sm.* 2011 demonstrates lower GSH-linked oxidation activity than the *Bradyrhizobium* (Fig. 4B). This may result from slight differences in the enzymes, leading to variations in turnover or sensitivity to assay conditions. Alternatively, the predicted ncRNA located between GSH-linked genes and *xoxF* in *Sm.* 2011 may influence regulation and result in less expression and activity of GSH-linked genes in *Sm.* 2011 compared to the *Bradyrhizobium* (Fig. 4A). FDH activity was also lower from *Sm.* 2011 than the *Bradyrhizobium* (Fig. 5). Sm2011_c04444 is the sole FdhA homolog found in *Sm.* 2011 and has 75% sequence similarity to Fdh*2*A from *Me.* AM1 and less than 40% similarity to other FDHs. Fdh2A from *Me.* AM1 is six-times less active in *in vitro* FDH assays compared to other FDHs present in *Me.* AM1 (19), explaining the low FDH activity from *Sm.* 2011. The *Bradyrhizobium* encode homologs of FDHs 1-4, consistent with their higher apparent FDH activity compared to *Sm.* 2011.

The proposed methanol assimilation pathway for *Sm*. 2011, *Bd.* 110, and *Bs.* 3456 is unlikely to be limited to these organisms alone. Instead, the XoxF-CBB pathway appears widespread among the *Bradyrhizobium*, *Sinorhizobium*, and *Mesorhizobium*. Selected members of each genus lack a complete serine/EMC cycle or RuMP pathway, but several encode a *xoxF* homolog as well as genes of the GSH-linked oxidation pathway and the CBB cycle (Fig. 7). This distribution implies that the capacity for Ln-dependent methanol assimilation using the CBB cycle may be more common in the rhizosphere than previously recognized. More work is needed to determine the conditions under which the XoxF-CBB pathway functions in the environment and how this pathway impacts microbial metabolism and influences plant health.

## Data Availability

Data is available from the corresponding author upon reasonable request. Transcriptomic data can be accessed via the GEO database (study number GSE295604) at: https://www.ncbi.nlm.nih.gov/geo/query/acc.cgi?acc=GSE295604.

## Acknowledgements

This material is based upon work supported by the National Science Foundation under Grant No. 2320667 and Grant No. 2127732. The authors acknowledge the BioAnalytical Facility at the University of North Texas for the support with mass spectrometry analyses during this work. This work used the Vincent J. Coates Proteomics/Mass Spectrometry Laboratory Core Facility, RRID:SCR_025852. We thank the Coates Lab at UC Berkeley for the use of their spectrophotometer.

---

**Table S1.**
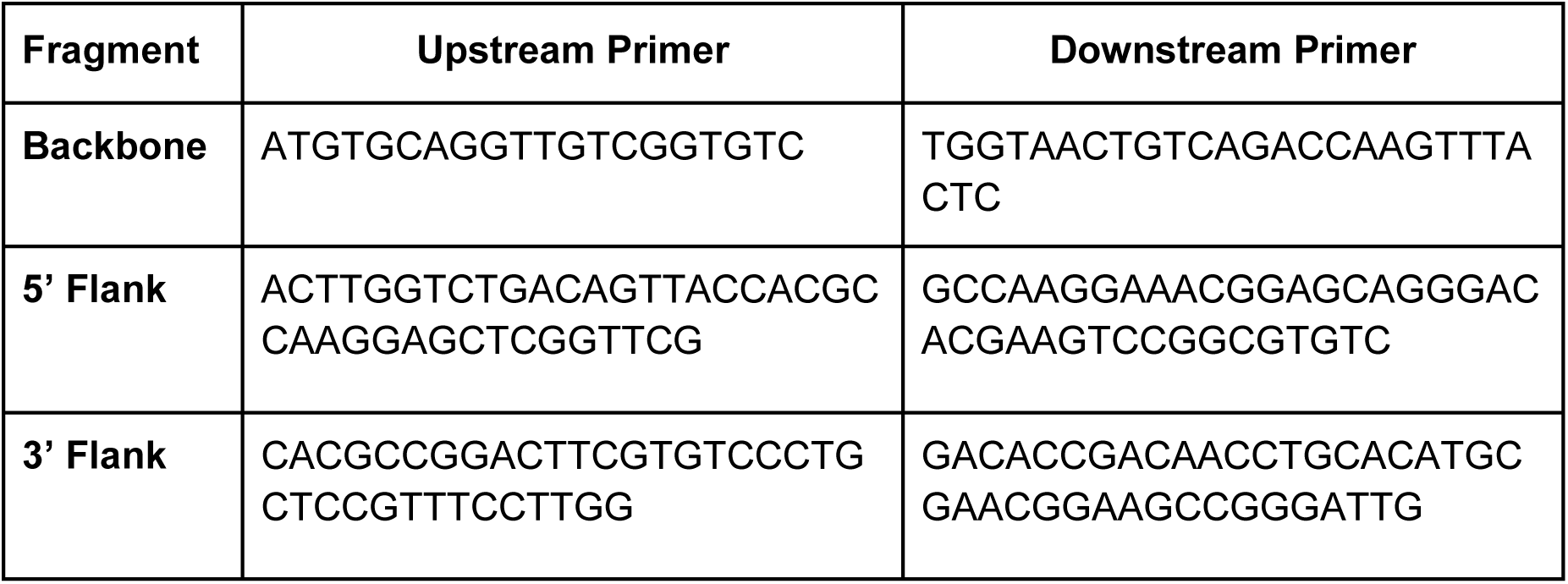
Primers used to generate plasmid pCRM031 used for the deletion of *cbbS* from *Bd.*110.

**Table S2.** Genes involved in Type I and Type II methylotrophy present in *Me.* AM1, *Bs.* 3456, *Bd.* 110, *Sm.* 2011. Genomes from MAGE were queried using EC number (when available) and via BLAST using the sequence of the gene of interest from *Me.* AM1 or *Bs.* 3456 when EC numbers were unavailable. The top hit for each gene is included in the table. In cases where multiple copies are present, only the first, by gene number, is shown for brevity. “-” indicates neither an annotated gene nor a homolog with greater than 50% sequence similarity could be identified.

| Gene | EC Number | *Me.* AM1 | *Bs.* 3456 | *Bd.* 110 | *Sm.* 2011 | Pathway |
| --- | --- | --- | --- | --- | --- | --- |
| *ftfL* | 6.3.4.3 | META1_0329 | - | - | SM2011_c02728 | H<sub>4</sub>F in *Me.* AM1 |
| *mtdA* | 1.5.1.- | META1_1728 | - | - | - | H<sub>4</sub>F in *Me.* AM1 |
| *fch* | 3.5.4.9 | META1_1729 | - | - | - | H<sub>4</sub>F in *Me.* AM1 |
| *purU* | 3.5.1.10 | - | VIDU01_420582 | AAV28_27315 | SM2011_c03205 | H<sub>4</sub>F Typical |
| *folD* | 1.5.1.5 | - | VIDU01_120226 | AAV28_04035 | SM2011_c02604 | H<sub>4</sub>F Typical |
| *mtdB* | 1.5.1.- | META1_1761 | - | - | - | H<sub>4</sub>MPT |
| *mch* | 3.5.4.27 | META1_1763 | - | - | - | H<sub>4</sub>MPT |
| *glyA* | 2.1.2.1 | META1_3384 | VIDU01_310356 | AAV28_22525 | SM2011_c01770 | Serine Cycle |
| *sga* | 2.6.1.45 | META1_1726 | VIDU01_170022 | AAV28_27730 | SM2011_a2139 | Serine Cycle |
| *ppc* | 4.1.1.31 | META1_1732 | VIDU01_10600 | AAV28_11795 | - | Serine Cycle |
| *mdh* | 1.1.1.37 | META1_1537 | VIDU01_270073 | AAV28_29905 | SM2011_c02479 | Serine Cycle |
| *mtkA* | 6.2.1.9 | META1_1730 | - | - | - | Serine Cycle |
| *mcl* | 4.1.3.24 | META1_1733 | - | - | - | Serine Cycle |
| *icl* | 4.1.3.1 | - | VIDU01_320092 | AAV28_09190 | SM2011_c00768 | Glyoxylate Shunt |
| *ms* | 2.3.3.9 | - | VIDU01_640301 | AAV28_04320 | SM2011_c02581 | Glyoxylate Shunt |
| *ccr* | 1.3.1.85 | META1_0178 | VIDU01_510004 | - | - | EMC Pathway |
| *pccA* | 6.4.1.3 | META1_3203 | VIDU01_310070 | AAV28_23715 | SM2011_b20756 | EMC Pathway |
| *mcd* | 1.3.99.10 | META1_2223 | - | - | - | EMC Pathway |
| *meaA* | 5.4.99.63 | META1_0180 | - | - | - | EMC Pathway |
| *mcMA* | 5.4.99.2 | META1_5251 | VIDU01_10717 | AAV28_12280 | SM2011_b20757 | EMC Pathway |
| *rpi* | 5.3.1.6 | META1_2299 | VIDU01_200722 | AAV28_15740 | SM2011_b20371 | RuMP Pathway |
| *hps* | 4.1.2.43 | - | - | - | - | RuMP Pathway |
| *phi* | 5.3.1.27 | - | - | - | - | RuMP Pathway |
| *glpx* | 3.1.3.11 | META1_0757 | VIDU01_10119 | AAV28_18895 | SM2011_b20202 | RuMP Pathway |
| *tkt* | 2.2.1.1 | META1_1861 | VIDU01_10121 | AAV28_04560 | SM2011_b20200 | RuMP Pathway |

**Table S3.** Key genes involved in the XoxF-CBB methanol assimilation pathway present in *Me.* AM1, *Bs.* 3456, *Bd.* 110, *Sm.* 2011. Genomes from MAGE were queried using EC number (when available) and via BLAST using the sequence of the gene of interest from *Me.* AM1 or *Bs.* 3456 when EC numbers were unavailable. The top hit for each gene is included in the table. In cases where multiple copies are present, only the first, by gene number, is shown for brevity. “-” indicates neither an annotated gene nor a homolog with greater than 50% sequence similarity could be identified.

| Gene | E.C. Number | *Me.* AM1 | *Bs.* 3456 | *Bd.* 110 | *Sm.* 2011 | Description of Gene Product Function |
| --- | --- | --- | --- | --- | --- | --- |
| *xoxF* | 1.1.2.10 | META1_1740 | VIDU01_860227 | AAV28_28625 | SM2011_b20173 | Ln-dependent PQQ methanol dehydrogenase. |
| *xoxG* | - | META1_1741 | VIDU01_860228 | AAV28_28630 | SM2011_b20174 | Cytochrome c. |
| *exaF* | 1.1.99.- | META1_1139 | VIDU01_860219 | AAV28_28595 | SM2011_b20173 | Ln-dependent PQQ ethanol dehydrogenase. |
| *gfa* | 4.4.1.22 | META1_3270 | VIDU01_860230 | AAV28_28640 | SM2011_b20186 | Glutathione-dependent formaldehyde activating enzyme. |
| *frmA* | 1.1.1.284 | - | VIDU01_860229 | AAV28_28635 | SM2011_b20170 | Glutathione-dependent formaldehyde dehydrogenase. |
| *frmB* | 3.1.2.12 | - | VIDU01_860197 | AAV28_28485 | SM2011_b20171 | S-Formyl Gluathione hydrolase. |
| *fdh1A* | 1.17.1.9 | META1_5032 | VIDU01_110101 | AAV28_08470 | - | W-Containing formate dehydrogenase. |
| *fdh2A* | 1.17.1.10 | META1_4848 | VIDU01_10810 | AAV28_12705 | SM2011_c04444 | Mo-Containing NAD-Dependent formate dehydrogenase. |
| *fdh3A* | 1.2.2.1 | META1_0303 | VIDU01_200818 | AAV28_24810 | - | Fe-S formate dehydrogenase. |
| *fdh4A* | - | META1_2094 | VIDU01_11139 | AAV28_40640 | - | Formate dehydrogenase. |
| *prk* | 2.7.1.19 | META1_0758 | VIDU01_10120 | AAV28_09870 | SM2011_b20201 | Phosphoribulokinase. |
| *cbbL* | 4.1.1.39 | - | VIDU01_10123 | AAV28_09885 | SM2011_b20198 | Large subunit of Rubisco. |
| *cbbS* | - | - | VIDU01_10124 | AAV28_09890 | SM2011_b20197 | Small subunit of Rubisco. |
| *cbbX* | - | - | VIDU01_10125 | AAV28_09895 | SM2011_b20196 | Rubisco regulator. |
| *lutA* | - | META1_1778 | VIDU01_860199 | AAV28_28495 | SM2011_b20178 | ABC transporter-periplasmic binding component. |
| *lutB* | - | META1_1779 | VIDU01_860200 | AAV28_28500 | SM2011_b20179 | Exported protein. |
| *lutE* | - | META1_1782 | VIDU01_860201 | AAV28_28505 | SM2011_b20184 | ABC transporter ATP-binding. |
| *lutF* | - | META1_1783 | VIDU01_860202 | AAV28_28510 | SM2011_b20185 | ABC transporter membrane component. |
| *lutG* | - | META1_1784 | VIDU01_860203 | AAV28_28515 | SM2011_b20169 | Exported protein. |
| *lutH* | - | META1_1785 | VIDU01_860194 | AAV28_28470 | - | TonB dependent receptor. |

**Fig. S1.**
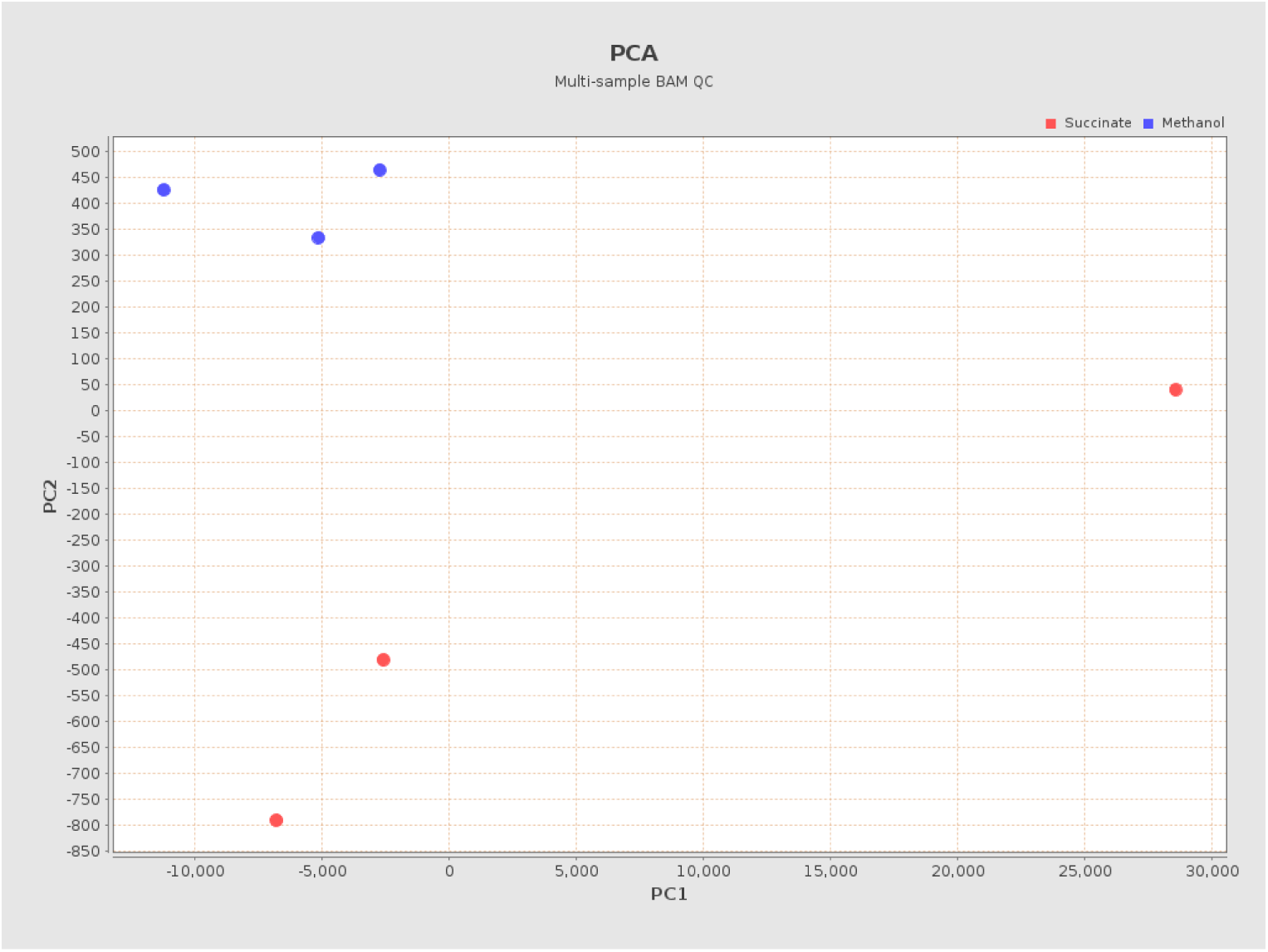
PCA plot of transcriptomic data from *Bs.* 3456 grown on succinate with La (red) and methanol with La (blue) generated from the HISAT2 - v2.1.0 module in KBase.

**Fig S2.**
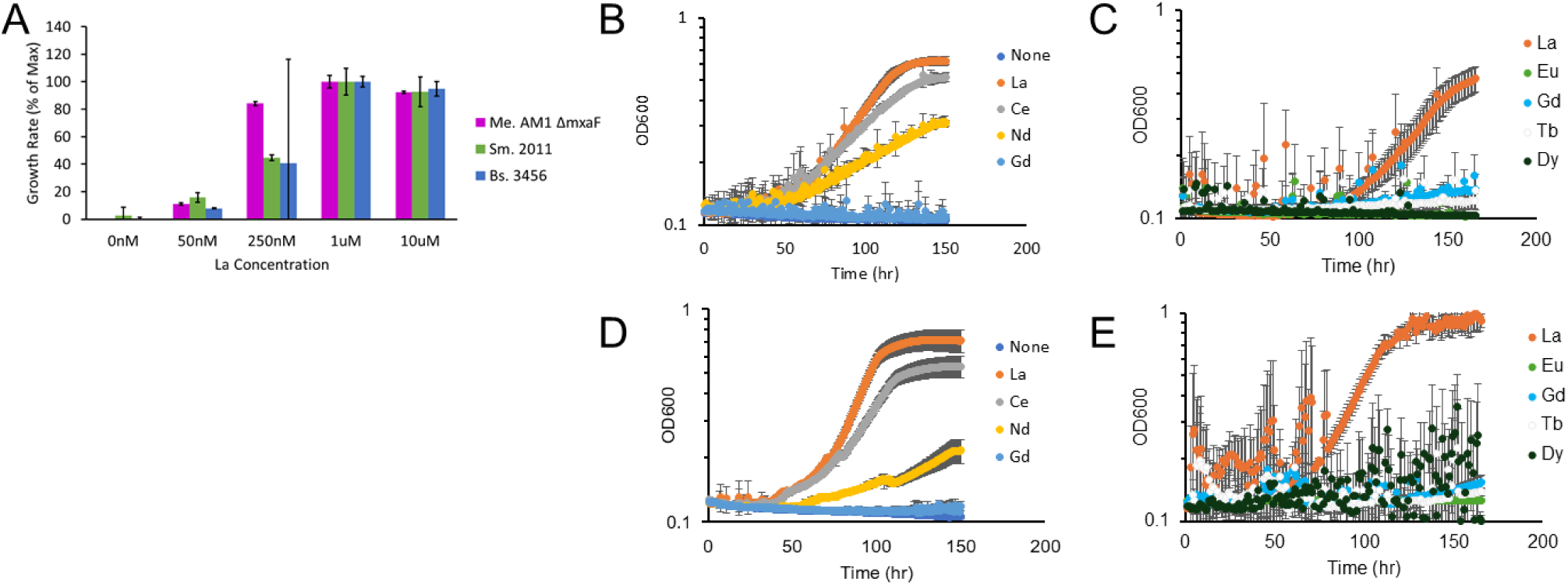
Growth of Bs. 3456 and Sm. 2011 on different concentrations and types of Lns. **(A)** Relative growth rates on 50mM methanol with increasing concentrations of LaCl_3_. According to a 2-tailed student’s t-test, for *Me.* AM1 growth rates in different concentrations of La were all different from one another at the p≤0.05 significance level. For *Sm.* 2011, all rates differed at the p≤0.05 significance level except for 1 vs 10 μM LaCl_3_. For *Bs.* 3456, only growth rates at 0 and 50 nM vs 1 and 10 μM were significantly different. **(B and C)** Growth of *Bs.* 3456 on 50 mM methanol with 10 μM of the indicated lanthanide source. **(D and E)** Growth of *Sm.* 2011 on 50 mM methanol with 10 μM of the indicated lanthanide source.

**Fig S3:**
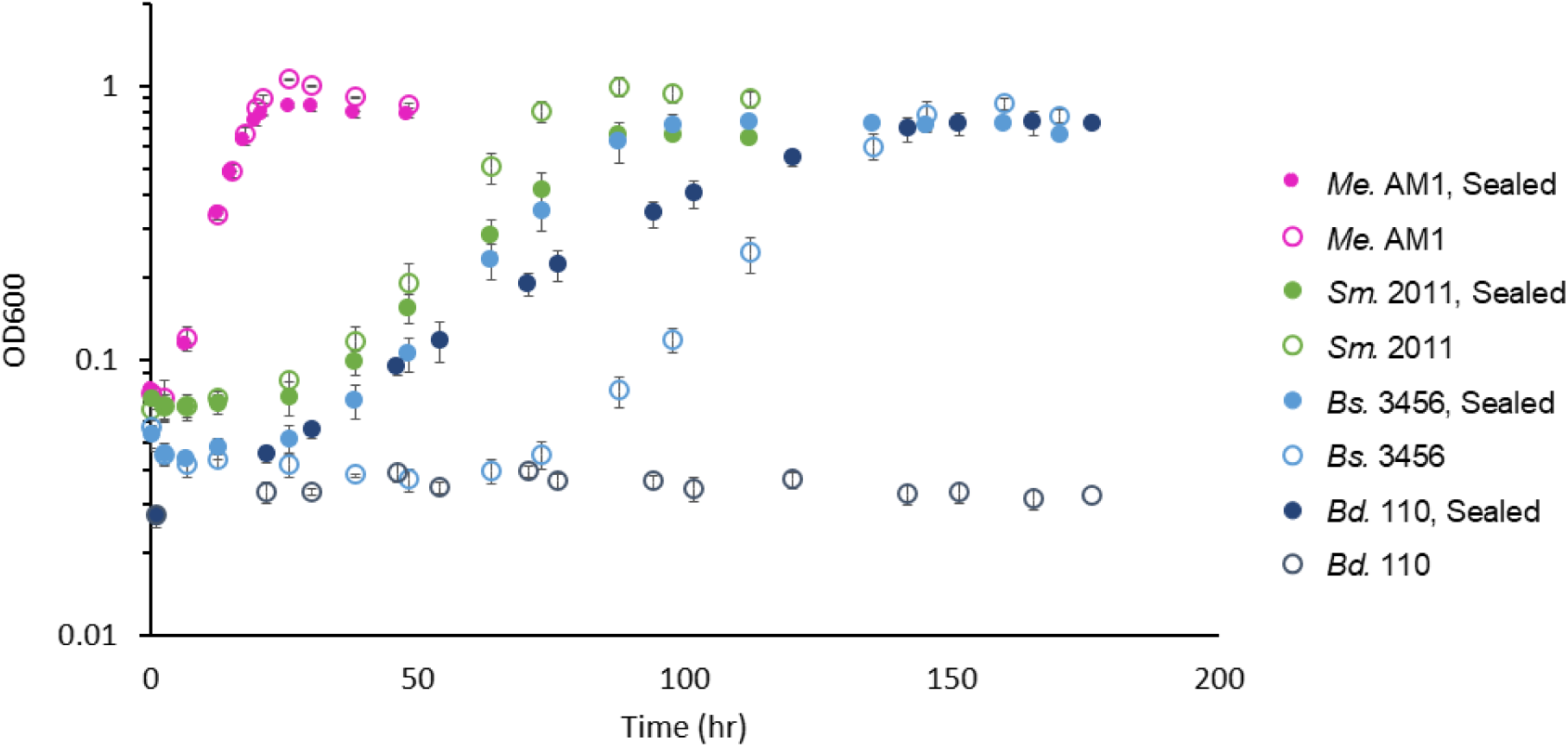
Growth in sealed (filled circle) vs foil-covered (open circle) Balch-style tubes facilitates growth of *Bd.* 110 and increases growth rate of *Bs.* 3456 on methanol. All samples were grown on 50 mM methanol with 10 μM LaCl_3_.

**Fig S4.**
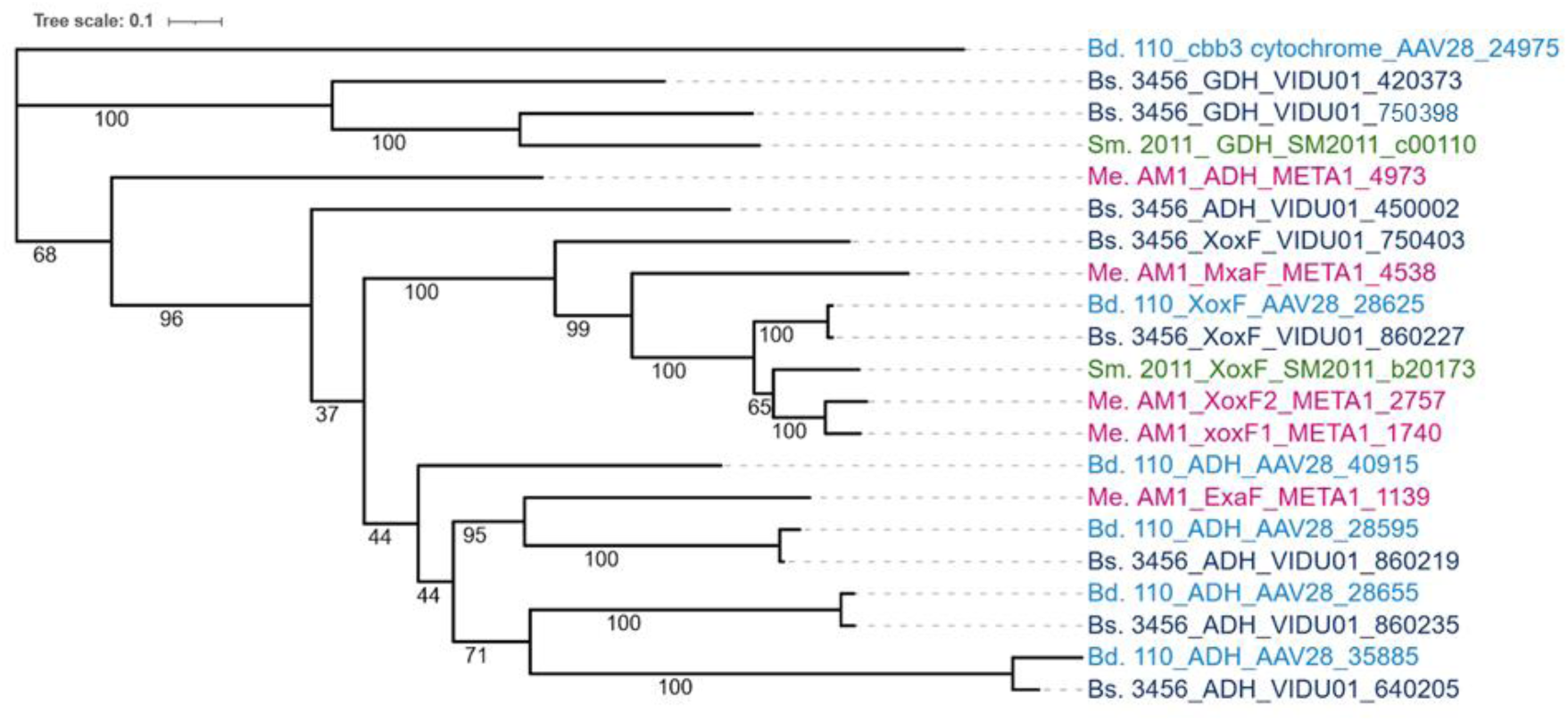
Phylogeny of Alcohol dehydrogenases found in the strains of interest. BLAST search using XoxF1 from AM1 as the query reveals multiple potential Ln-dependent proteins encoded in each genome of interest. Strain names, gene product annotations, and accession numbers are provided and color-coded according to the strain. *Bd.* 110 light blue, *Bs.* 3456 dark blue, *Sm.2011* green, *Me.* AM1 pink. All but AAV28_24975, META1_4973, META1_4538, and VIDU01_450002 are predicted to be Ln-dependent based on the presence of an ‘additional Ln-binding Asp’ in the active site.

